# High-efficiency HDR in immortalized cell lines by crude rAAV mediated donor template delivery

**DOI:** 10.1101/2022.05.02.490359

**Authors:** Stuti Mehta, Altantsetseg Buyanbat, Ge Zheng, Nan Liu, Stuart H. Orkin

## Abstract

Owing to low efficiency of homology-directed repair (HDR), precise knock-in (KI) of large DNA fragments is a challenge in genome editing. High-efficiency HDR has been reported for primary cells in preclinical gene therapy by combining CRISPR/Cas9 mediated induction of double-strand breaks (DSB) with delivery of a single-stranded DNA HDR-donor-template via highly purified recombinant adeno-associated virus (rAAV). Due in part to the labor and expense of rAAV particle purification, rAAV-mediated HDR-template delivery has been underutilized used to generate large KIs in cultured cell lines. Here, we report application of *crude* preparations of rAAV to deliver HDR-templates for the KI of large ∼2kb fragments at various genomic loci in several -human as well as mouse cell lines at high efficiency. Our approach should facilitate experiments necessitating KI of large DNA fragments to tag endogenous loci for visualization and/or conditional protein degradation.

## Introduction

Protein function is often deduced on the basis of cellular localization, protein partners, and genetic loss of function phenotypes. Antibodies that detect a protein of interest (POI) are critical tools for these studies, but may not be available or be of sufficient quality for experimental studies. Epitope (or other) tagging of POIs is an alternate and exceedingly useful approach, which is optimally executed by knock-in of DNA sequences to generate protein fusions at N-or C-termini. Endogenous tagging circumvents limitations associated with exogenous transgene expression by preserving native promoter activity and protein function. It also facilitates visualization of proteins for cellular localization and/or trafficking, and chromatin occupancy studies for those chromatin-associated factors for which specific antibodies are unavailable. Moreover, tagging with a variety of degrons has enabled targeted, conditional and reversible degradation of the fused POI, providing an unprecedented opportunity for kinetic analysis of downstream effects of protein loss, as opposed to steady-state measurements following genetic ablation [1-3] [4-6].

Electroporation of a ribonucleoprotein (RNP) complex of Cas9 nuclease and a single synthetic guide RNA (sgRNA) into cells yields high-efficiency genetic editing and few off-targets owing to a short *in vivo* half-life of RNPs [7, 8]. Upon provision of a donor template in trans, precise endogenous KIs can be generated by homology-directed repair (HDR) of the CRISPR/Cas9 induced DSBs [9, 10]. Methods of donor delivery include transfection of single-or double-stranded oligonucleotides, plasmid DNA, or transduction with HDR-template carrying replication-deficient lentivirus, or recombinant adeno-associated virus (rAAV) [11, 12]. Concerns regarding cellular toxicity associated with transfection of large amounts of naked DNA limit the availability of large HDR-donor molecules in the nucleus and adversely affect KI efficiency [13, 14]. To circumvent these issues, HDR templates have been introduced with viral vectors, such as AAV.

AAV is a non-pathogenic, replication-deficient, ssDNA virus of the *parvoviridae* family [15]. Its linear, single-stranded, ∼4.7 kb genome flanked by 147 bp inverted terminal repeats (ITRs) can be replaced entirely by recombinant DNA of similar size to generate recombinant AAV (rAAV, Figure 1A) [15, 16]. Upon transduction, rAAV delivers the recombinant DNA cargo into the host nucleus. The cargo does not integrate into the host cell genome and instead persists as episomes that are diluted out with successive cell divisions [17]. Transduction with several natural, or synthetic serotypes of AAV engineered to carry HDR templates have been used to achieve high-efficiency HDR following electroporation of a Cas9/sgRNA [18] [19]. However, AAV purification protocols are notoriously laborious and expensive. As such, save a few reports in permissive cell lines, rAAV has been underutilized for KI of large DNA fragments into cell lines [20-24]. Here, we dispense with the extensive purification of AAV. Our experiments show that unpurified, crude rAAV preparations used to deliver ∼1.2 kb HDR-templates consistently achieve high-efficiency HDR in a variety of cell lines, and at distinct genomic loci. Our approach to KI generation is inexpensive, highly efficient, versatile, and facilitates genetic modification of cells for many experimental purposes.

**Figure 1.**
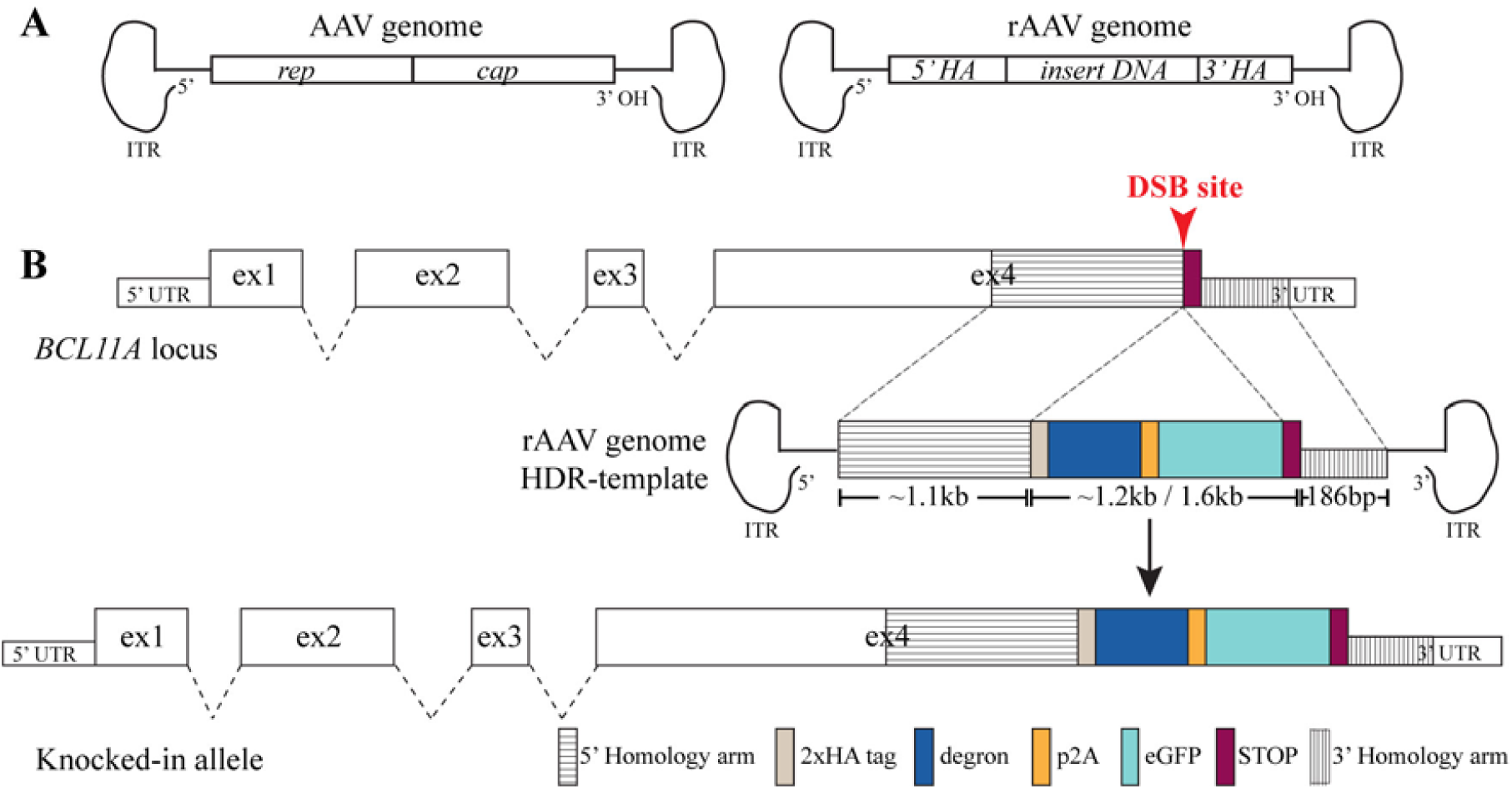
Engineering of the AAV genome to generate rAAV carrying an HDR donor template. ***A***, Left, Illustration of the single stranded ∼4.7 kb AAV genome flanked by two ITR sequences. Right, A recombinant AAV genome. The entire ∼4.7 kb region between the ITRs is replaced with recombinant HDR-donor-template to generate rAAV. ***B***, Design of the HDR-template to KI either the FKBP12^F36V^ (∼1.2 kb) or SMASh (∼1.6 kb) degrons at the C-terminus of *BCL11A*. Homology arms are indicated by striped boxes. Red arrow indicates site of double strand break (DSB). Figure is not to scale.

## Results

### Experimental design

BCL11A is a zinc-finger transcription factor that is a critical repressor of γ-globin genes in adult erythrocytes [25-28], and also important for B-lymphocyte and nervous system development [29, 30]. To enable functional studies, we set out to tag the endogenous *BCL11A* gene in immortalized human CD34-like progenitor cells, HUDEP-2. We generated three BCL11A fusions: two degrons: dTAG and SMASh, and the ALFA-tag to degrade BCL11A via the Trim-away technology. The dTAG system consists of a degron termed FKBP12^F36V^ that selectively binds one valency of a bivalent molecule, dTAG-47, which additionally recruits CRBN -a ubiquitously expressed E3 ubiquitin ligase and a member of the ubiquitin-proteasome protein degradation, inducing rapid proteolysis of the fusion protein [6]. The SMASh (Small-Molecule Assisted Shut-off of protein function) system is composed of two components: the HCV-protease NS3 and a destabilizing degron. In the absence of a small molecule inhibitor (Asunaprevir), the protease cleaves itself and the degron from the fusion protein partner, rendering the POI in its native form. Treatment with Asunaprevir leads to retention of the degron, and destabilization, followed by lysosomal/proteosomal degradation of the fused protein [4]. To generate fusions of BCL11A with FKBP12^F36V^, SMASh, or ALFA-tag, we first determined that HUDEP-2 cells are diploid for the *BCL11A* locus (data not shown). The degrons were placed at the C-terminal of BCL11A-XL (Figure 1B) [31, 32].

### Order of RNP delivery and rAAV2/6 transduction in HUDEP-2 cells

We tested the effect of the order of RNP delivery and rAAV transduction on HDR efficiency. rAAV transduction prior to delivery of Cas9/sgRNA RNP might be anticipated to ensure immediate availability of the donor template upon the formation of a DSB. However, in a recent report, nucleofection enhanced rAAV6 transduction as well as KI efficiency in primary CD34^+^ cells [19]. We observed that transduction of HUDEP-2 cells with rAAV2/6 carrying a GFP gene, closely following RNP nucleofection (N-AV, ∼20 min) similarly resulted in increased transduction (Figure S2B), indel efficiency (p<0.05) (Figure 2A), as well as HDR (Figure 2B) compared to the delivery of RNP 24h after viral transduction (AV-N). HUDEP-2 cells were transduced either before or after RNP nucleofection at a relative volume of 1:4 [= crude viral prep:media volume]. The physical titer of the crude viral lysates was ∼6×10^6^ vgs/cell. For all subsequent experiments, rAAV transduction closely followed RNP nucleofection (∼20min).

**Figure 2.**
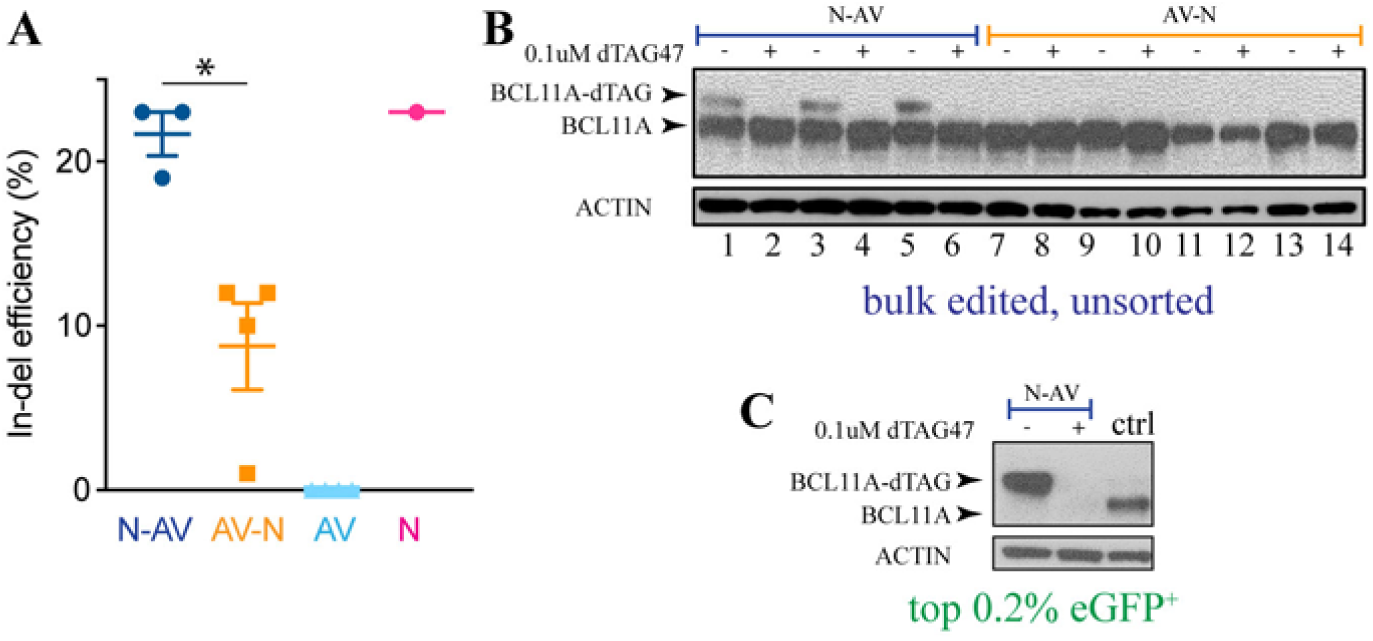
High-efficiency KI of ∼1.2 kb insert containing the FKBP12^F36V^ degron at the *BCL11A* locus by Cas9/sgRNA RNP nucleofection followed by crude rAAV2/6 transduction of the HDR-donor-template. ***A***, Efficiency (%) of indel formation when RNP nucleofection precedes crude rAAV2/6 transduction (N-AV) and vice-versa (AV-N), as compared to only RNP nucleofection (N), or only crude rAAV2/6 transduction (AV). Unpaired students *t* test, n=3, error bars represent *s*.*e*.*m*. Western blot for BCL11A in ***B***, bulk edited HUDEP-2 cells, and ***C***, the 0.2% highest eGFP^+^ expressing cells, with either N-AV or AV-N delivery and 15h treatment with 0.1μM dTAG-47 (+) or DMSO control (-), *n* = 3-4. ACTIN is the loading control.

### High-efficiency KI of large DNA fragments at the BCL11A locus in HUDEP-2 cells

We assessed modification of the endogenous BCL11A locus 7 days after RNP nucleofection and FKBP12^F36V^ HDR-donor containing rAAV2/6 transduction. The HDR cassette contained homology arms of 778bp and 186bp flanking 1212bp FKBP12^F36V^-p2a-eGFP. By Western blot with antibody to BCL11A, we observed a ∼14 kDa super-shift of BCL11A in the edited bulk cells, consistent with fusion of FKBP12^F36V^ to BCL11A (hereafter, BCL11A-FKBP12^F36V^, Figure 2B, lanes 1,3,5). PCR screening of single-cell clones from the edited pool revealed that ∼15% and 4% clones contained heterozygous or homozygous KI alleles, respectively (Figure S2B, Table 1). KI was in-frame with *BCL11A* in all clones tested (Figure S2D).

**Table 1:**
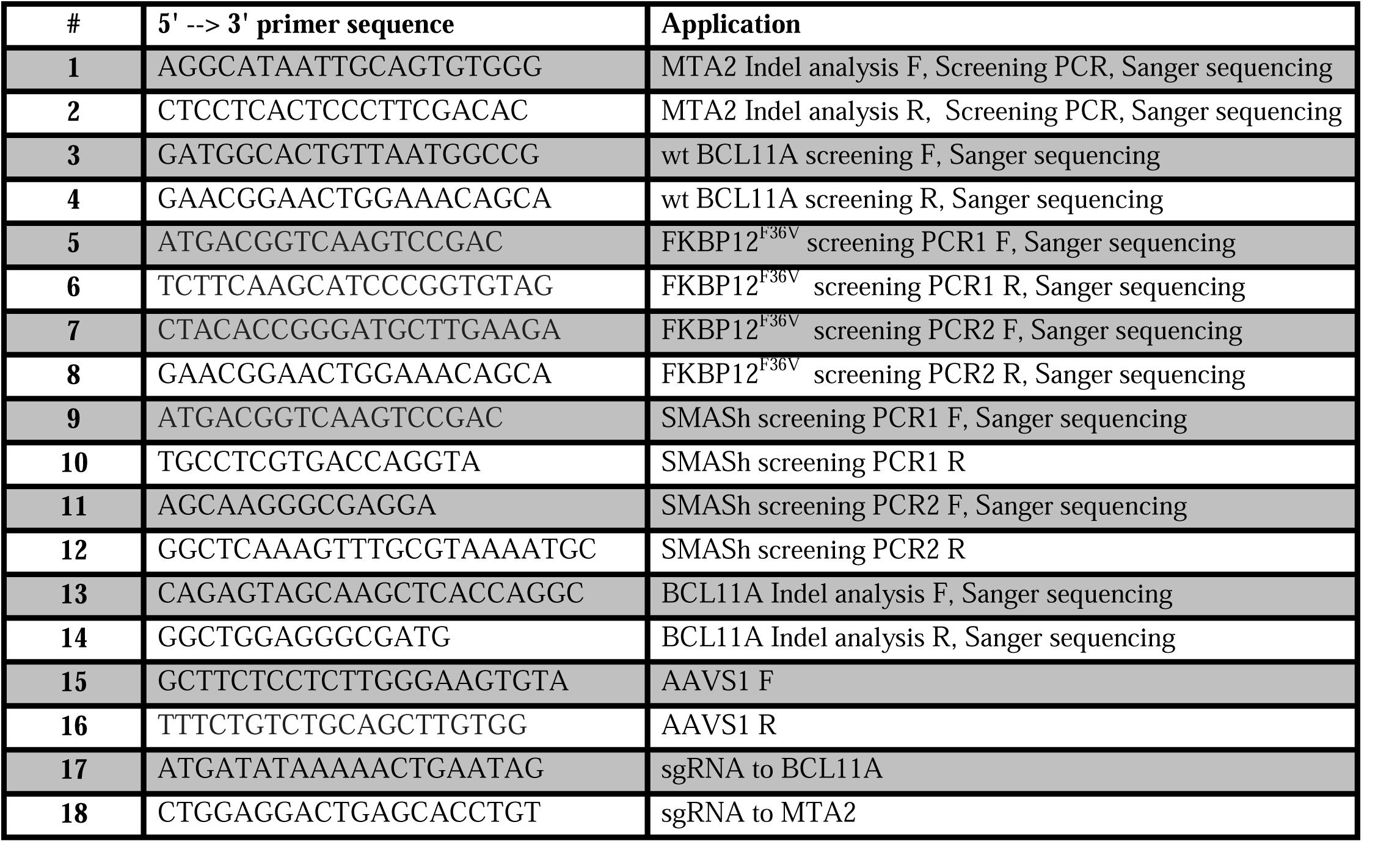
Sequence information for primers and sgRNA used in this study.

About ∼0.2% of the highest eGFP expressing cells were collected by FAC-Sorting of the edited pool (Figure S2C). Wildtype-sized BCL11A was not detected on Western blot, whereas a band consistent with BCL11A-FKBP12^F36V^ was detected, and subsequently lost upon treatment with dTAG-47 (0.1 μM, 15 h, Figure 2C). PCR and Western blot based screening revealed ∼90% of the sorted pool harbored homozygous KI, whereas a minor fraction had heterozygous knocked-in alleles or no insertions. One monoallelic KI clone showed NHEJ mediated ∼19 bp deletion at the 3’ end of the other *BCL11A* allele (Figure S2D).

Nucleofection of Cas9/sgRNA RNP followed by crude-rAAV2/6 mediated transduction of SMASh HDR-donor-template resulted in an allele KI frequency of ∼22% as determined by PCR genotyping of single-cell derived clones. About 7% of clones were targeted at both alleles, whereas ∼ 30% clones were tagged at a single allele.

We obtained a similar efficiency of KI of xxbp ALFA-tag at the C-terminal of BCL11A. Talk about how we used this tag to degrade BCL11A with the trim-away system.

### Crude rAAV2/6 mediated HDR at the MTA2 locus in HUDEP-2 cells

We tested applicability of this method at other genomic loci by inserting in tandem, one V5 and two HA tags followed by P2A-eGFP (total ∼0.9 kb) immediately preceding the stop codon at the C-terminus of a NuRD subunit MTA2 [33, 34]. Seven days after RNP and crude rAAV2/6 delivery, a distinct population, ∼14.5% of the cells, expressed GFP^+^, suggesting a high rate of in-frame incorporation of the HDR-template (Figure 3C). The top ∼50% of the GFP^+^ population was collected and limiting dilution performed to obtain single-cell clones. PCR screening of 182 individual GFP^+^ clones with primers annealing to genomic regions outside the homology arms (Figure 3B, Table 1) indicated that ∼59%, ∼35%, and 4% of clones harbored heterozygous, homozygous, or no KIs, respectively. Aberrant KI was observed in 18% of clones, where a PCR product whose size was inconsistent with wildtype or knocked-in alleles was observed (Figure 3B). Precise homozygous KI was confirmed in 30 clones by Sanger sequencing. Western blots of two representative homozygous KI clones with an MTA2 and an HA antibody show a super-shift in the size of MTA2 consistent with the desired KI (Figure 3D).

**Figure 3.**
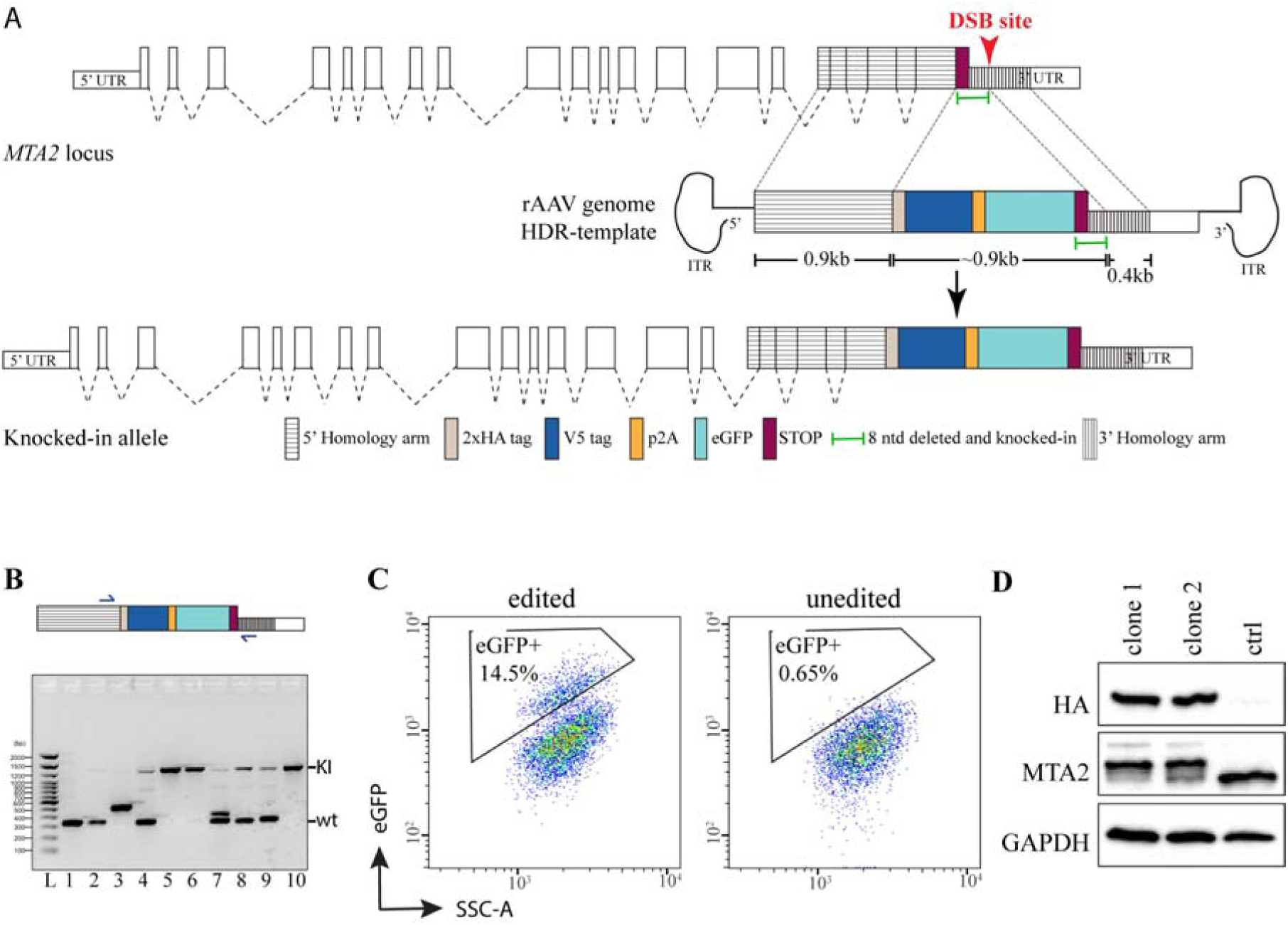
Knock-in of ∼0.9 kb DNA containing the V5 tag, 2xHA and eGFP at the *MTA2* locus by Cas9/sgRNA RNP nucleofection followed by delivery of HDR-template via crude rAAV2/6 preparations. ***A*, Top:** Illustration of the *MTA2* genomic locus, Middle: the design of the HDR-template, and Bottom: the KI allele. Figure is not to scale. ***B***, PCR screening strategy for the *MTA2* KI. *Top*, Approximate sites of primer annealing (indicated by small arrowheads) at the *MTA2* locus. *Bottom*, Representative agarose gel electrophoresis image of PCR results for various *MTA2* genotypes: wildtype (wt, lane 1,2), homozygous KI (lane 5,6,10), heterozygous KI (lane 4,8,9) and aberrant KI (lane 3,7). Lane L is a DNA ladder. ***C***, FACS plots showing expression of eGFP^+^ cells in the edited population (left) compared to unedited, control HUDEP-2 cells (right). ***D***, Western blot of two individual eGFP^+^ clones carrying homozygous KI at the *MTA2* locus, as well as unedited HUDEP-2 cells as control, with antibody to HA as well as to MTA2. GAPDH is the loading control.

### Precise KI of large DNA fragments at the BCL11A locus in HCC1187 cell line

To test application of this approach to other cell lines, we sought to knock-in the FKBP12^F36V^-2xHA-p2A-eGFP tag at the C-terminus of *BCL11A* in the immortalized breast cancer line HCC1187 [35]. We first screened the infection efficiency of 8, premade GFP-expressing AAV serotypes, and determined that AAV2 showed the highest efficiency of transduction in HCC1187 cells (Figure 4A). Nucleofection of Cas9/sgRNA RNP was followed by transduction of crude rAAV2/2 harboring the FKBP12^F36V^-HDR-donor-template. After 10 days, 0.5% of the highest eGFP expressing cells were collected by FACS. Western blotting for BCL11A detected only a super-shifted “BCL11A-FKBP12^F36V^” in this population, which was undetectable after treatment with dTAG-47 (0.2 μM or 0.5 μM, 24 h, (Figure 4B). Sanger sequencing confirmed precise in-frame KI of FKBP12^F36V^ in this population.

**Figure 4.**
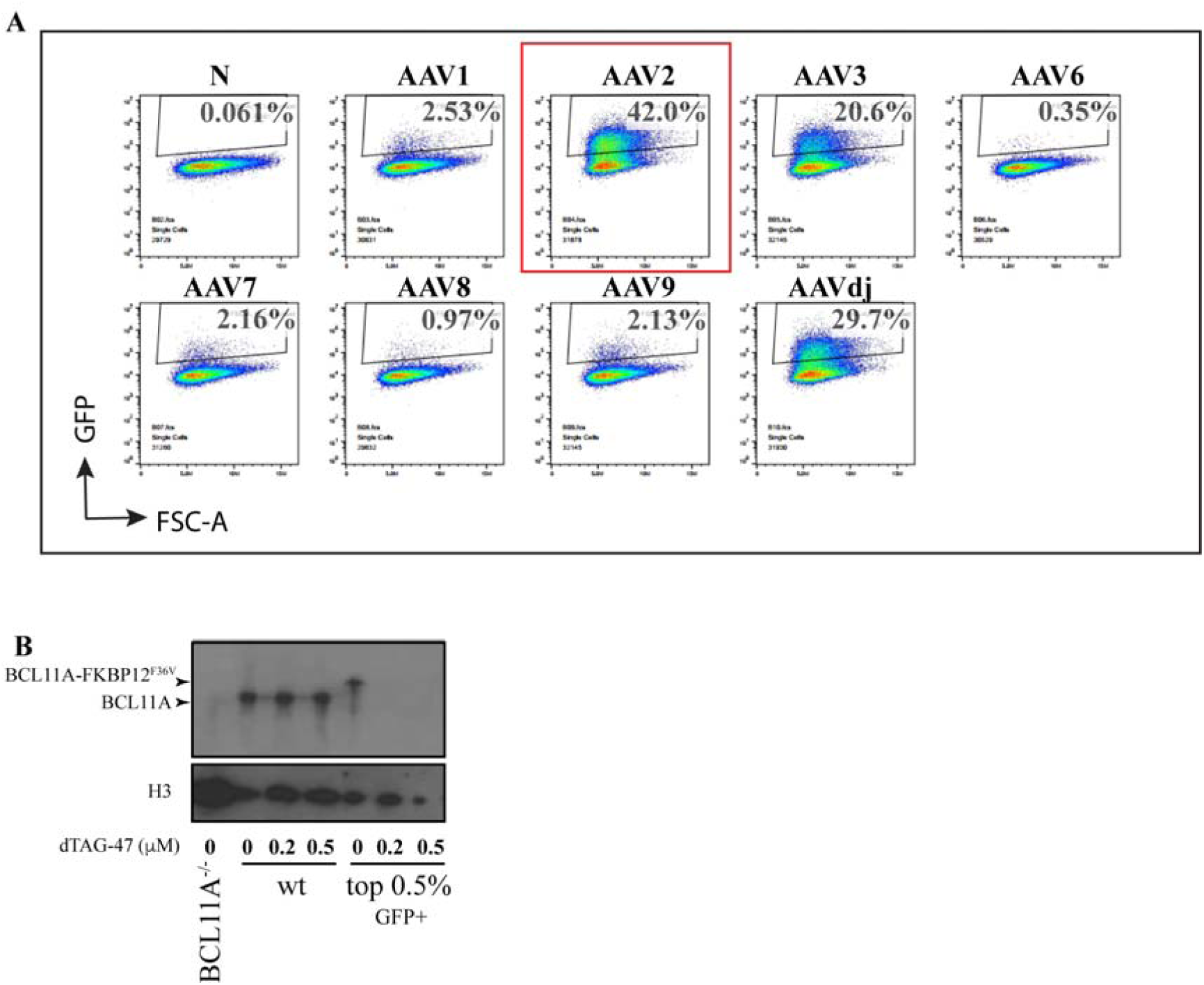
Knock-in of ∼1.2 kb DNA containing the FKBP12^F36V^ degron at *BCL11A* in HCC1187 cells by RNP nucleofection followed by delivery of HDR-template via crude rAAV2/2 preparations. ***A***, FACS plots showing infection efficiency of 8 unique AAV viral serotypes in HCC1187 cells, as measured by % of GFP^+^ cells. ***B***, Western blot for BCL11A in *BCL11A*^*-/-*^ HUDEP-2 cells, unedited HCC1187 cells and the 0.5% highest eGFP^+^ expressing edited HCC1187 cells, after 24 h treatment with 0.2 μM dTAG-47, 0.5 μM dTAG-47 or DMSO treatment control (0 μM dTAG-47). H3 is the loading control.

## Material and Methods

### Cell culture

HUDEP-2 cells were cultured at a density of 5×10^4^-5×10^5^/mL in expansion medium as described elsehere [36]. Briefly, StemSpan Serum-Free Expansion Medium (SFEM, StemCell Technologies) was supplemented with 2 × 10^2^ IU/mL Penicillin-Streptomycin, 50 ng/mL recombinant human stem cell factor (R&D systems), 3 IU/mL Epoetin-alfa (Amgen), 0.4 μg/mL dexamethasone (Sigma-Aldrich) and 1 μg/mL doxycycline (Sigma-Aldrich). The day before RNP electroporation, cells were diluted to at a density of 7.5 × 10^3^/mL with the expansion medium.

HCC1187 cells were cultured in RPMI-1740 with 10% heat-inactivated fetal bovine serum and 1% Penicillin-Streptomycin. Cells were passaged every 3-4 days and cell density maintained between 0.5-1.5 × 10^6^/mL.

### Testing the infection efficiency of 8 AAV serotypes in HCC1187 cell line

AAVPrime™ Adeno-associated virus (AAV) Serotype Testing Kit (GeneCopoeia) was used to determine the efficiency of 8 different AAV serotypes in infecting the HCC1187 cell line. Each AAV serotype in this kit encapsidates an identical genome, containing AAV2 terminal ITRs flanking a CAG-GFP expression cassette. Briefly, ∼1.3 × 10^5^ cells were plated in 10 wells of a 24-well plate in 1 mL HCC1187 culture medium. At 80% confluence, the media was replaced with 200 μL of media containing 2 µl AAV particles (one serotype per well) and the plate incubated for 2 hours at 37 °C, with swirling every 30 minutes. One well received no virus and was used as a negative control. Thereafter, 500 µL fresh medium was added to each well. After 24 h, cells were harvested and percent infection by each serotype was assessed by flow cytometry.

### sgRNA selection and design of the HDR-donor cassettes

BCL11A knock-ins: Four spCas9 guide RNAs (NGG PAM), all yielding∼25% indel efficiency are available in the ∼40 bp region proximal to the stop codon of BCL11A-XL (IDT CRISPR-Cas9 guide RNA design checker, data not shown). An sgRNA that would introduce a DSB immediately 5’ of the stop codon was selected to enable the 5’ and the 3’ homology-arms of the HDR donor template to be adjoining the DSB (Figure 1B, Table 1). The close proximity of homology arms to DSB was expected to enhance the efficiency of precise KI [37]. The 5’ and 3’ homology arms were ∼1.1 kb and 186 bp in size respectively. A single nucleotide, silent mutation was introduced to the stop codon contained in the 3’ homology arm to prevent cleavage of the HDR-template or re-cleavage of the locus by Cas9 nuclease following HDR [37]. The homology arms flanked FKBP12^F36V^-2x-P2A-eGFP or SMASh-2x-P2A-eGFP, bringing the total length of inserts to 1.2 kb for FKBP1212^F36V^ and 1.6 kb for SMASh (Figure 1B).

MTA2 knock-in: A ∼53% efficient sgRNA with a cut-site 5 bp upstream of the stop codon was chosen (Table 1). A 400 bp homology arm adjoined 3’ of the DSB site, whereas a 900 bp, 5’ homology arm was located 8 ntd upstream (Figure 3A). The missing 8 ntd include the stop codon and 5 ntd of the 3’UTR; and were included in the donor-template, to be re-inserted upon HDR. No silent mutations were introduced in this region to preserve the integrity of the regulatory 3’ UTR (green feature in Figure 3A).

### Assembly and screening of pAAV2 transfer plasmids

Three ug of pAAV2 vector (Addgene #89771) was digested with MluI/BstEII to release the insert between the two ITRs. The HDR donor was cloned between two AAV2 ITRs as three overlapping dsDNA oligonucleotides (gBlocks, IDT, shown as red, blue, yellow in Figure S1A) with NEBuilder HiFi DNA assembly at 50°C for 1 hour at manufacturer recommended molar ratios. One Shot Top10 chemically competent *E. coli* were transformed with 2 μL of the completed HiFi reaction. Clones carrying the desired recombinant HDR-template were identified by Sanger sequencing. The palindromic AAV ITRs form secondary hairpin structures hindering Sanger sequencing through these regions. ITRs also frequently recombine in bacterial host environments, resulting in deletion of one or both ITRs, and subsequent failure of AAV packaging [38]. To confirm the integrity of the ITRs candidate plasmid clones were digested overnight with *Sma*I, which cuts twice within the recombination-prone region of the ITR, but not in the vector backbone. Intact plasmids yield a backbone band of ∼2.6 kb, as well as insert-specific bands. For the FKBP12^F36V^ donor, additional bands of ∼1.5 kb, ∼500 bp, ∼480 bp, ∼280 bp, ∼80 bp are expected (Figure S1B). Similarly, the MTA2 donor was assembled into pAAV2 using two gblocks. Correctly assembled constructs were used as transfer vectors for packaging into rAAV2/6 or rAAV2/2 (see below).

### Preparation of rAAV2/6 and rAAV2/2 crude lysates

The triple-transfection method was used to generate crude rAAV lysates [39]. HEK293T cells were plated in a 10 cm dish in 1X Penicillin-streptomycin-Glutamine and 10% FBS in DMEM. At 80% confluence, the medium was replaced with fresh medium. Then, cells were triple-transfected using polyethylenimine Max (PEI Max, Polysciences 24765-1) with 12 μg of pAAV-helper, 7.5 μg of pRep2Cap6, and 7.5 μg of transfer plasmid (PEI:DNA = 3:1), in DMEM without phenol red (Life Technologies 31053036). 3 days after transfection, cells were scraped and collected by spinning at 1300 RPM for 5 min. Cell pellet was re-suspended in 1-2 mL DPBS without calcium or magnesium and lysed by three rounds of freeze-thaw. This was accomplished by placing them alternately in a dry ice/ethanol bath until completely frozen and in a water bath of 37 °C until completely thawed. After the final thaw, cell lysate was spun at 1300 RPM for 5 min and the supernatant was filtered through a 0.22 μm syringe filter to yield a crude viral lysate. Viral lysates were stored in 25 μL aliquots at 4°C for up to 4 weeks, and at -80 °C thereafter.

### rAAV2/6 titration by qPCR

The physical titer of the crude rAAV preps was obtained by qPCR with primers specific to the AAV2 ITR sequence which detect both ssDNA derived from rAAV6/2 viral particles as well as dsDNA of a pAAV2 transfer plasmid [40]. Briefly, a control plasmid containing the FKBP12^F36V^ insert was purified from a bacterial clone (QIAGEN Plasmid Maxi kit). Six serial dilutions of the plasmid were generated, spanning a range of 2 × 10^8^-2 × 10^3^ plasmid molecules/ μL. In parallel, the crude rAAV2/6 lysates were treated with DNase I (ThermoFisher) at 37°C for 30 min to eliminate contaminating, extra-viral plasmid DNA. Six dilutions of the DNase-treated crude viral lysate (10x, 1000x, 5000x, 25000x, 125000x) were made in PBS. qPCR was then performed in a 20 μL reaction volume with 2×SYBR Green qPCR mix (Qiagen), 100 nM each of the sense and antisense primers (Table 1), and 5 μL of serially diluted template DNA (either the plasmid standard or the crude AAV prep), in duplicate. The PCR reaction was as follows: 98°C for 3 min; 40x [98°C for 15 sec / 58°C for 30 sec]; melt curve. The physical titer of viral preps [viral genomes (vgs)/μL] was calculated based on the standard curve and sample dilutions.

### RNP nucleofection and crude rAAV2/6 transduction in HUDEP-2 cells

HUDEP-2 cells in the exponential phase of growth (< 2 × 10^5^/mL) were used. Cas9/sgRNA RNP was generated by mixing 120 pM guide RNA (2’-O-methyl analog and 3’ phosphorothioate internucleotide modified-sgRNA, custom ordered from Synthego Corp.) with 61 pM Cas9-Alt-R protein (IDT) and incubated at room temperature for 10 min. Meanwhile, 5 × 10^4^ HUDEP-2 cells were collected by centrifugation at 500xg for 5 min, washed in 1X room temperature PBS, and re-suspended in 20 μL NF-solution (P3 Primary Cell 4D-Nucleofector™ X Kit, Lonza). Thereafter, RNP was mixed with the cells, and nucleofection carried out in a 4D-nucleofector X Unit with program EO-100. Immediately after nucleofection, cells were supplemented with 80 μL warm expansion medium. After 10 min, cells were collected by centrifugation at 500xg for 5 min and re-suspended in 100 μL expansion medium in one well of a flat-bottom 96-well plate. At this point, 20 μL crude rAAV viral prep was added to the cells, and the cells returned to the 37°C incubator for 3 days. When the cells were transduced by rAAV2/6 before nucleofection, 20 μL of crude rAAV2/6 lysate was added to 5×10^4^ cells/100 μL of media. After 24 h, 50K cells were nucleofected as described above and returned to the incubator in 100 μL expansion medium in one well of a flat bottom 96-well plate, and left to recover for 3 days. The two experiments were analyzed in parallel.

### RNP nucleofection and rAAV2/2 transduction in HCC1187 cells

To prepare RNP, 120 pmol modified sgRNA (Synthego corp) was mixed with 100 pmol Cas9 nuclease (Alt-R IDT) and incubated at RT for 10 min. Meanwhile, one million HCC1187 cells were trypsinized, collected by centrifugation at 500xg for 5 min and resuspended in SF solution (SF cell line 4D-Nucleofector™ X Kit, Lonza). After addition of the RNP, cells were nucleofected with program CM120. Cells were collected by centrifugation and transferred to 1 mL of culture medium. For rAAV2/2 mediated knock-in, 150 μL of crude rAAV2/2 was added to the cells within 30 min of nucleofection. Fifty thousand cells were collected after 3 days for PCR analysis, and the rest were cultured for 7 days until sorting.

### Assessment of Indel efficiency

Genomic DNA was extracted using Quickextract solution (Lucigen). A ∼700 bp-1 kb region flanking the sgRNA annealing site was amplified by PCR 5-7 days after RNP nucleofection. Cas9/Scrambled sgRNA was the negative control. The efficiency of sgRNAs to induce indels was measured by comparing Sanger sequence traces between the targeting sgRNA vs control samples using either the ICE (Synthego Corp) or TIDE analysis freeware.

### Generation of HUDEP-2 monoclonal cell lines and colony screening

About 4-7 days after RNP and rAAV delivery, or after FAC-Sorting, monoclonal cell lines were generated by limiting dilution cloning of the edited cells. Based on hemocytometer counts, cells were diluted to a density of ∼30 cells/10 mL and seeded into round-bottom 96-well plates with a multichannel micropipetor, resulting in roughly 30 of the 96 wells containing single cells. In about 24 h after cell seeding, most cells center to the bottom of the round-bottom wells and are easily visualized under an upright light microscope. At this stage, any wells containing more than 1 cells were flagged and later excluded from the analysis to minimize the possibility of generating mixed clones. Clones were supplemented with 50 μL fresh expansion medium after 5 days in culture, and at 10 days, all media was replaced with fresh 100 μL expansion medium per well. After 15 days in culture, DNA was extracted from the clones by cell lysis with Quickextract DNA extraction solution (Lucigen), and used as template for PCR with 2x Hotstart PCR mix (Qiagen) and 100 nM of forward and reverse primers each. Each 96-well plate was screened with 3 PCRs: 1) Positive control gDNA: PCR with primers that annealed to the universal AAVS region 2) PCR to detect a wild type *BCL11A* allele, i.e. that had not undergone HDR 3) PCR with primers that specifically amplify a knocked-in allele. Frequently, for long inserts, the third PCR was split into two – PCR1 amplified the 5’ region of a KI allele, and PCR 2 amplified the 3’ region (Indicated by Pink and dark blue arrowheads in Figure S2B). A combination of steps 2 and 3 enabled distinguishment between homo/heterozygous KIs. Primer sequences are in Table 1.

### Western Blot

Cell pellets were re-suspended in RIPA buffer (Boston Bioproducts) supplemented with a protease inhibitor cocktail (Roche), and incubated on ice for 15 min to generate whole cell lysates. Lysates were clarified by centrifugation at 20,000 x g at 4°C for 10 min. Identical number of cells from each sample was collected and electrophoresed on polyacrylamide gels (Bio-Rad). Antibodies used were as follows: BCL11A (Rabbit monoclonal cat# ab191401, Abcam, 1:3000 dilution), β-actin C4 (cat# sc-47778 HRP, Santa Cruz Biotechnology, 1:5000 dilution), GAPDH (Rabbit monoclonal lot 14C10, cat#2118S, Cell Signaling Technology, 1:1000 dilution), HA (12CA5, Mouse mAb, cat# 11583816001, Roche, at 1 μg/mL final concentration); MTA2 (Rabbit polyclonal antibody, cat# ab8106, Abcam, 1:1000 dilution), H3 (cat# ab24834, Abcam, 1:5000 dilution). Imaging was performed by detection of HRP conjugated primary/secondary antibody on an x-ray developer.

### Fluorescence Activated Cell Sorting

Flow cytometry was used to collect cells that had undergone HDR. Cells expressing the highest eGFP were sorted into one well of a 96-well plate containing fresh expansion medium/RPMI, using a FACS Aria II SORP (BD Biosciences).

## Discussion

Here we describe a simplified approach for generation of endogenous knock-in modifications of genes of interest in cell lines leveraging Cas9/sgRNA RNP nucleofection followed by delivery of the HDR donor template via crude rAAV preparations. The approach reduces the time to generation of stable KI clones in two ways: 1) elimination of extensive rAAV purification procedures and 2) generation of high-efficiency, precise KI, which reduces the need for extensive screening of single-cell clones for transgenic lines. We have piloted the strategy in HUDEP-2 cells, a platform that facilitates genetic analysis of globin gene regulation, as well as in HCC1187 cells.

We observed higher transduction rates, significantly more indel formation and increased HDR when rAAV mediated HDR delivery followed RNP nucleofection (N-AV), as compared to rAAV transduction preceeding RNP nucleofection (AV-N). AAV transduction proceeds through several steps. First, virus is internalized by endocytosis [41]. In CD34^+^ cells, nucleofection increased endocytotic capacity of the host cells, leading to an increase in viral transduction by rAAV [19]. A similar mechanism is likely to operate in the, CD34^+^-like HUDEP-2 cells used in our study. The efficiency of KI is partially dependent on the intracellular availability of the HDR template [13, 14]. Consistent with this notion, we observed higher HDR in N-AV compared to AV-N. HDR in N-AV is likely additionally augmented by the higher indel frequency in N-AV compared to AV-N.

As with other methods of donor delivery, the size of HDR donor template affects KI efficiency. In HUDEP-2 cells, KI of the SMASh tag which was ∼1.6 kb in size (in addition to ∼1.3 kb homology arms), was substantially less efficient compared to that of FKBP12^F36V^ KI, which was ∼1.2 kb in size, or the ∼0.9 kb KI at the *MTA2* locus. For integration of larger gene cassettes, the size-dependence can be overcome by enrichment of KI alleles with fluorescence or antibiotic-resistance marker selection. As illustrated here, it is feasible to enrich for a nearly pure population (>90%) of KI alleles by fluorescent-marker selection. In cell lines for which derivation of stable clonal lines is challenging, a population of cells enriched for the KI may suffice. To enrich specifically for homozygous KI clones, double selection has been used [42, 43]. Two unique selection-carrying HDR-templates may be delivered simultaneously by transduction with two crude rAAV preps, or by generating a heterogeneous rAAV preparation by supplying the two transfer plasmids to one well of a packaging cell line. Similarly, multiplex transductions may be used to target multiple loci in a cell line.

In contrast to ultrapure rAAV, crude viral preps contain impurities, such as cell debris, empty capsids or AAV-encapsidated DNA impurities. While it is critical to minimize such contaminants in clinical settings, this may be unnecessary for routine experimental purposes [44]. Of relevance to *in vitro* applications, endotoxin carried over from the plasmids used for the generation of crude lysates may induce cytotoxicity in the host cell line, and can be avoided by use of endotoxin-free plasmid DNA isolation methods. Indeed, we did not observe excess cell death in our transductions compared to nucleofection-only controls.

A disadvantage of rAAV for DNA delivery is the size of its cargo capacity, which is limited to ∼4.7 kb. Considering homology arms are of 400 bp each, KIs of upto ∼3.9 kb can be generated [45]. Finally, integration-deficient rAAV (lacking the wildtype AAV genome) may randomly integrate into the genome at a frequency of 0.1-1% [17, 46]. Save for genome screening to detect random integration events, we recommend analysis of several independent knock-in clones in order to attribute a phenotype to a POI/locus with confidence.

For application of this method in cell lines other than those described here, several parameters need to be examined. An electroporation protocol that yields maximal indel efficiency while ensuring minimal cell death may need optimization. In addition, the appropriate AAV serotype for a given cell line should be identified. Lastly, crude rAAV preparations are routinely generated for pilot or trial experiments, and can be custom ordered from commercial vendors or core facilities. With the discovery and advent of various Cas9 nucleases, different PAM specificities may be leveraged [47-50]. Application of the methods we have described to novel Cas9s will also expand the possibilities of precise KI in cell lines and facilitate diverse experimental objectives.

---

**Figure S1.**
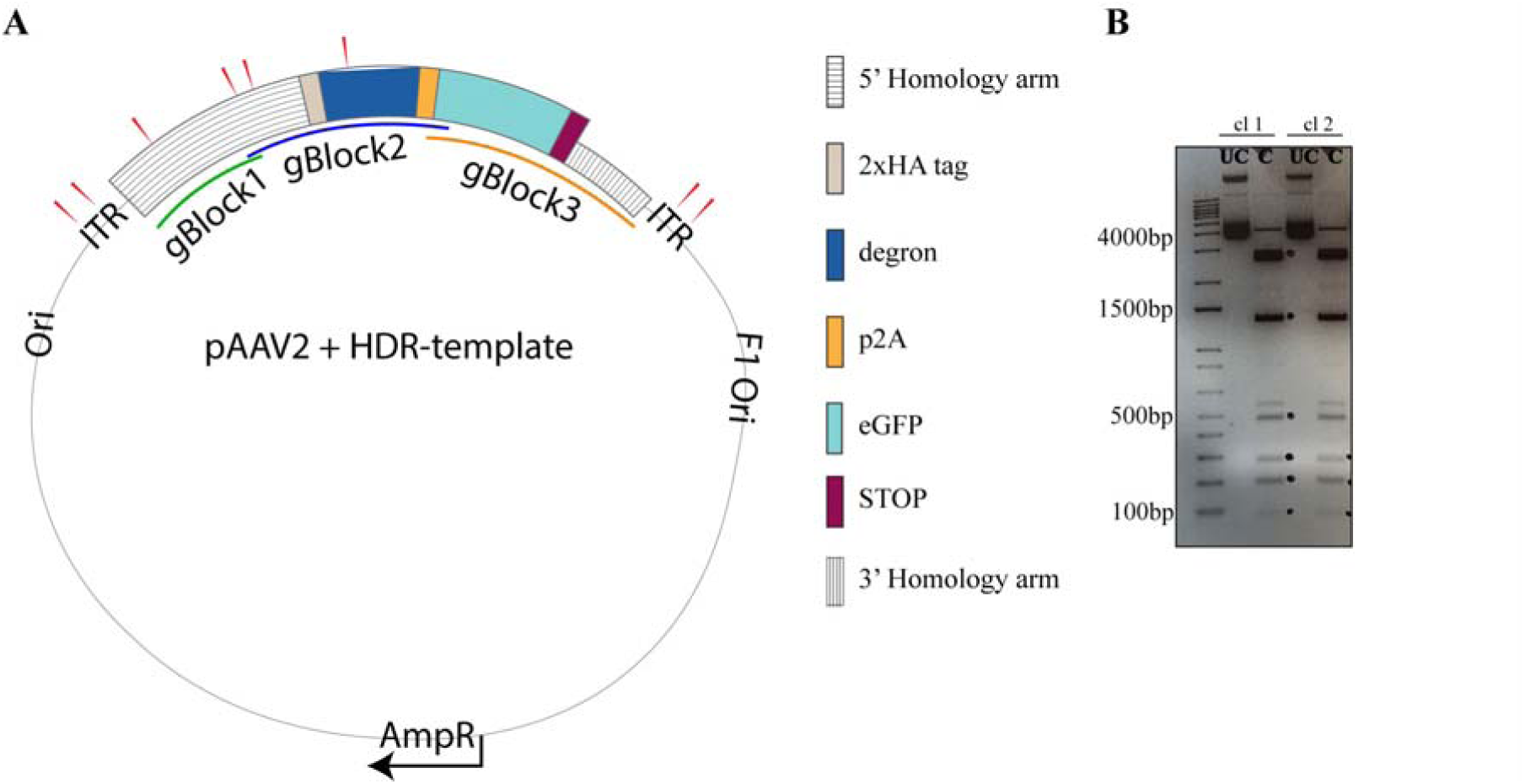
Construction of the pAAV2 transfer plasmid. ***A***, Diagram of the recombinant pAAV2 transfer-plasmid construct containing the HDR template for KI of degron sequences, between the two ITRs. The green, blue and orange lines represent overlapping gBlocks used for NEBuilder HiFi DNA assembly into the pAAV2 plasmid. Red pointers indicate approximate sites for *Sma*I digestion. Figure is not to scale. ***B***, Agarose gel electrophoresis image of *Sma*I digest (C) for two pAAV2 clones (labeled cl1 and cl2). Uncut (UC) plasmids are loaded alongside as controls. Restriction fragments of expected lengths are marked on the image with black dots. Plasmids from clones cl1 and cl2 are intact (ITRs not recombined), and can be used for packaging into rAAV6.

**Figure S2.**
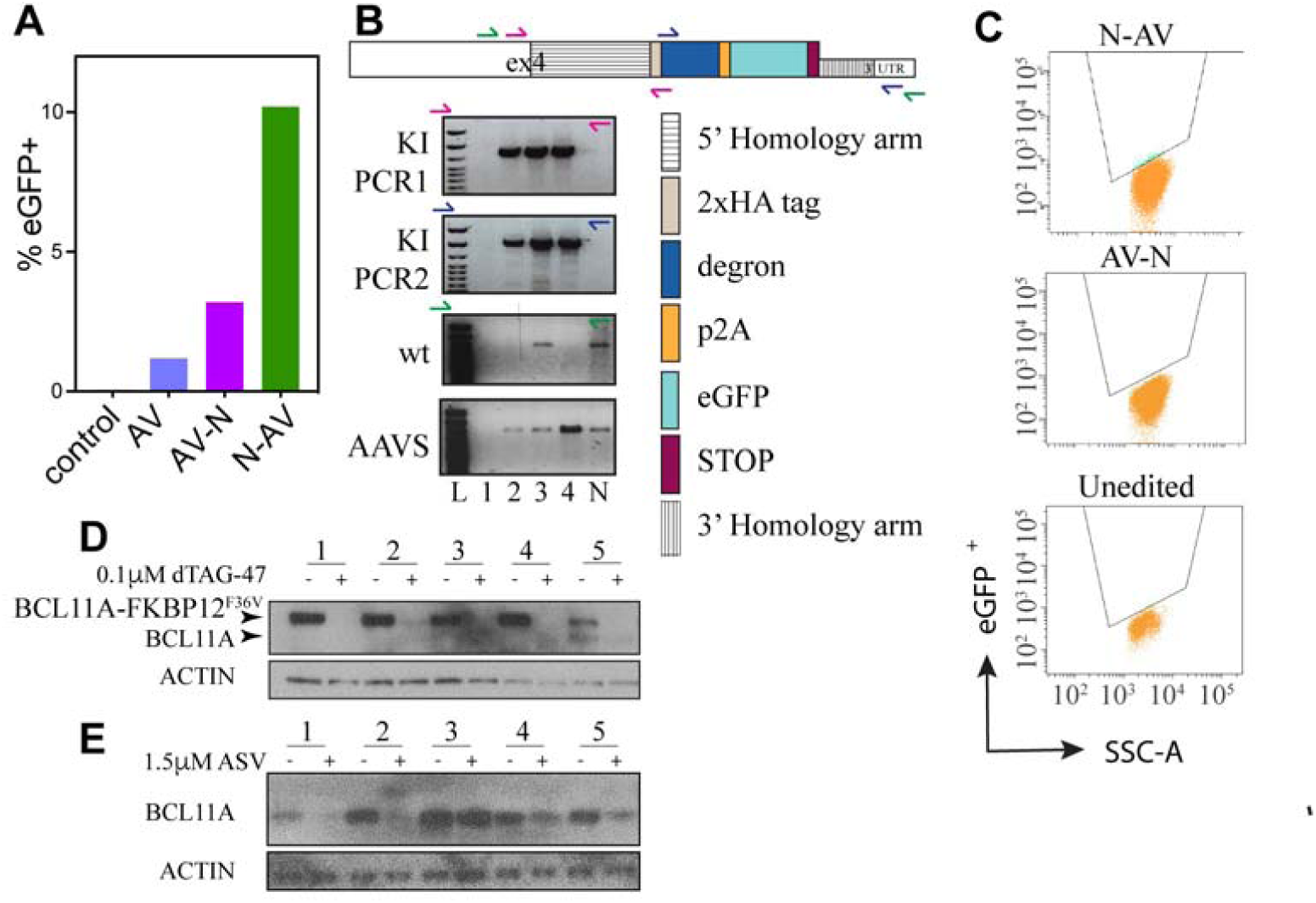
***A***, Percentage of eGFP^+^ HUDEP-2 cells after ∼7 days when RNP nucleofection with a non-targeting sgRNA precedes transduction with crude rAAV2/6 expressing GFP under a CAG promoter (N-AV) and vice-versa (AV-N), as compared to only crude rAAV2/6 transduction (AV), and unedited HUDEP-2 cells (control). ***B***, PCR screening strategy for the two degrons systems. *Top*, Approximate sites of primer annealing (indicated by small arrowheads) at the wildtype or KI *BCL11A* locus. Primer pairs are color-matched. Figure is not to scale. *Bottom*, Representative agarose gel electrophoresis image of PCR results for various *BCL11A* genotypes with respect to FKBP12^F36V^ KI: wildtype (lane 5), homozygous KI (lane 2,4), heterozygous KI (lane 3) and no template control (lane 1). Lane L is a DNA ladder. Primers to the AAVS region served as a positive control for PCR. ***C***, FACS plots showing expression of eGFP^+^ cells in the N-AV population (top) compared to AV-N (middle), and unedited HUDEP-2 cells (bottom). ***D***, Representative Western blot of screening strategy for FKBP12^F36V^ KI clones. Individual clones were treated with 0.1 μM dTAG-47 (+) or DMSO control (-) for 15 h. Five clones are shown, lanes 1,2 and 4 show complete degradation of BCL11A-FKBP12^F36V^, indicating homozygous KI. Lane 3 is a clone that has a monoallelic FKBP12^F36V^ KI with an out of frame deletion of the stop codon. Lane 5 shows a heterozygous KI clone, with the wildtype band unaffected upon treatment with dTAG-47. ACTIN is the loading control. ***E***, Representative Western blot of the screening strategy for SMASh KI in individual clones. Single cell clones derived from a pool of eGFP^+^ cells were treated with 1.5 μM ASV (+) or DMSO control (-) for 48 h. Five clones are shown, lanes 1 and 2, homozygous KI clones show near complete degradation of BCL11A. Lane 3, a clone without KI shows no degradation upon ASV treatment, whereas lane 4 and 5 show diminished BCL11A indicating presence of heterozygous KI. ACTIN is the loading control.

## Acknowledgements

We thank the BCH viral core for preparing crude viral lysates, the DFCI FACS core facility for helping with cell sorting, Behnam Nabet for discussions on employing the dTAG strategy and providing the dTAG-47 reagent, Divya Vinjhamur and Daniel Bauer for sharing *BCL11A*^*-/-*^ HUDEP-2 cells. S.H.O. is an Investigator of the Howard Hughes Medical Institute, and supported by NHLBI R01HL032259 and P01HL032262.

## Author contributions

SM conceived and led the study under the supervision of SHO. SM designed and performed the KIs at *BCL11A* in HUDEP-2 cells. AB provided technical support. GZ designed and performed KI at *MTA2* in HUDEP-2 cells, NL performed the KI at *BCL11A* in HCC1187 cells. SRM and SHO wrote the manuscript with input from all authors.

### Abbreviations used

HDR: Homology directed repair
rAAV: recombinant Adeno-Associated virus
KI: Knock-In
DSB: Double strand break
POI: Protein of interest
RNP: Ribonucleoprotein
sgRNA: Single guide RNA
ITR: Inverted terminal repeats
SMASh: Small molecule assisted shut down of protein function
ASV: Asunaprevir
N-AV: Nucleofection followed by AAV transduction
AV-N: AAV transduction followed by Nucleofection
CAG: CMV Early Enhancer/Chicken Beta Actin (CAG) Promoter
PBS: Phosphate buffered saline

