## Supplementary figures and images for "High-efficiency HDR in immortalized cell lines by crude rAAV mediated donor template delivery"

### Supplemental figure S1

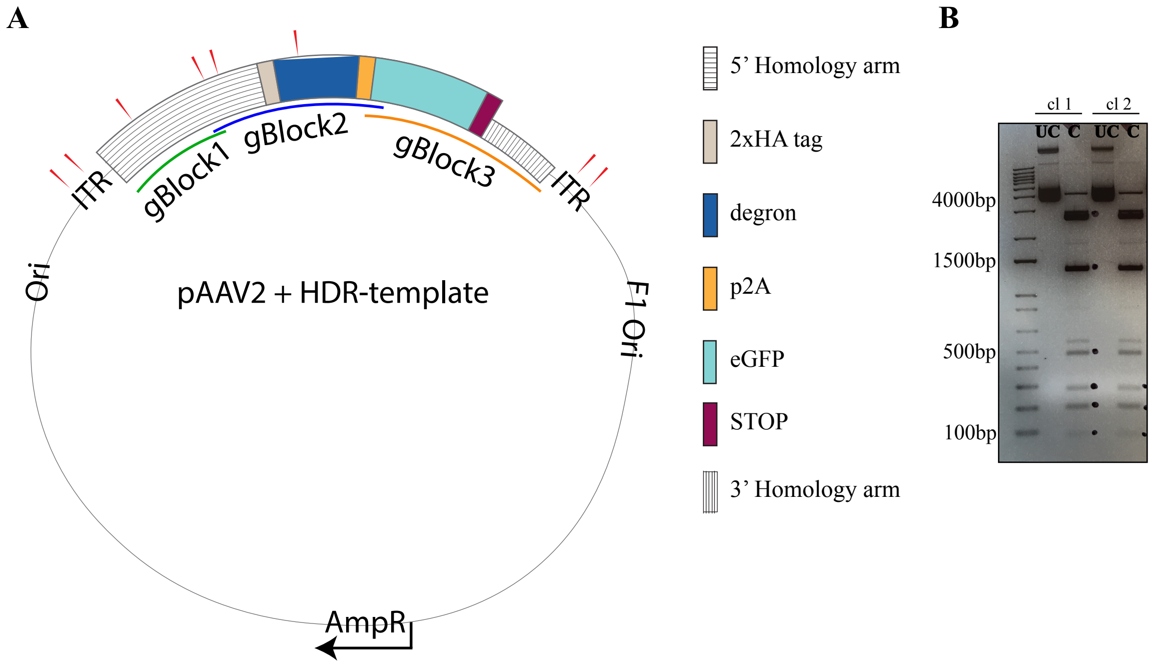

### Supplemental figure S2

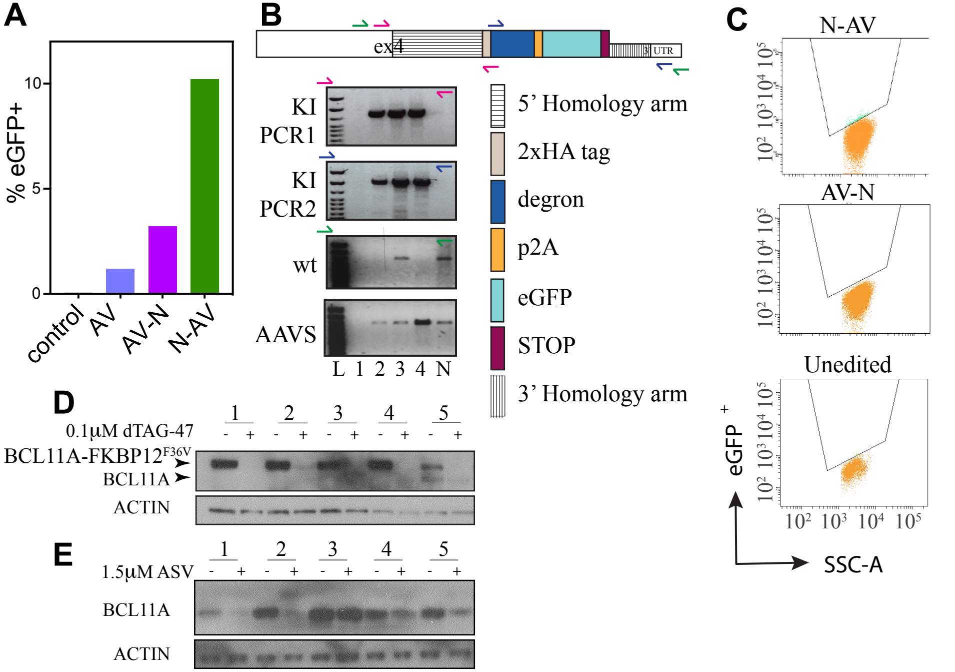
